# Cetuximab produced from a goat mammary gland expression system is equally efficacious as innovator cetuximab in model systems of cancer

**DOI:** 10.1101/2020.06.04.135434

**Authors:** Qian Wang, William Gavin, Nick Masiello, Khanh B. Tran, Götz Laible, Peter R Shepherd

## Abstract

Humanised monoclonal antibodies have proven a very effective mode of therapy for a wide range of conditions. With many of the monoclonal antibody drugs now coming off patent there is an increasing interest in developing biosimilar, or even biobetter, forms of these drugs. With the commercial competition associated with such generic products there is increasing demand for improved production and purification system for biosimilars. Cetuximab, also known as ‘Erbitux’, is widely used in cancer therapy, especially in advanced colorecal cancer, metastatic non-small cell lung cancer and head and neck cancer and it will come off patent in the near future. We have previously reported on a genetically engineered goat system to produce cetuximab (gCetuximab) in milk. Herein we now report on the further charactization of the gCetuximab produced utilizing additional and more sophisticated biological assays. There is similar bioactivity of the gCetuximab compared with the commercial product produced in mammalian cell culture. In particular both cetuximab antibodies selectively target EGFR and induce its internalization to down regulate EGFR signaling. Both forms have very similar half life in animals and in a HT29 colorectal cancer xenograft model have similar efficacy. We also show that the toxin MMAE can be conjugated to gCetuximab, that this targets it to cells and that this results in direct cell killing in HT29 cells. This demonstrates that the gCetuximab will also be a viable vehicle for antibody drug conjugate based therapies. Taken together, this shows that the goat milk monoclonal antibody production system is an effective way of producing a biosimilar form of cetuximab.

## Introduction

The monoclonal antibody based cancer treatment approach has proven to be one of the most successful cancer therapeutic classes of drugs over the last two decades due to their high target-specificity to tumour cells and low cytotoxicity (1). Cetuximab, an anti-EGFR monoclonal antibody, is obtained by attaching the Fv variable regions of a murine monoclonal anti-EGFR antibody (C225) to human IgG1 heavy and kappa light chain constant regions (2). Known by its trade name Erbitux, cetuximab is an epidermal growth factor receptor (EGFR) inhibitor which is used for cancer treatment, especially in advanced colorectal cancer, metastatic non-small cell lung cancer as well as head and neck cancer (3). Cetuximab is a monoclonal antibody of the immunoglobulin G1 (IgG1) subclass, which binds to EGFR with high affinity, thereby blocking the endogenous EGFR ligands from binding. This results in inhibition of the function of the EGFR receptor (4–6). Cetuximab is selective and does not bind to other HER family receptors, such as ErbB2, ErbB3 and ErbB4. EGFR is constitutively expressed in many normal epithelial tissues. Its signaling pathways are involved in the regulation of cell survival, cell cycle progression, angiogenesis, cell migration and cellular metastasis (6–9). Over-expression of EGFR is frequently detected in many human cancers. Cetuximab was approved by the the US Food and Drug Administration (FDA) in February 2004 and by the European Medicines Agency (EMA) in June 2004 as a cancer therapy that used alone or in combination with other medications to treat colon or rectal cancer that has spread to other parts of the body (2).

Clinical grade cetuximab (Erbitux), is produced by expressing it in expensive mammalian cell bioreactors, driving interest in alternative production systems. We have recently reported the development of a system for expressing cetuximab in the mammary gland of lactating transgenic goats and purifying this version of cetuximab (gCetuximab) from the resulting milk (10). Our earlier report described the gCetuximab and provided evidence that it may be a “Biobetter”as; (a) it has unique characteristics and benefits that increase its safety profile (i.e. no α-gal linkage or chance for immunogenicity), and (b) it has an increased antibody dependent cellcular cytotoxity (ADCC) profile (10). This report describes a range of studies that demonstrate that the biological properties of the cetuximab produced in the transgenic milk system are equivalent to those of the commercial cell culture product. Together this provides evidence that the goat produced product could be taken forward as a potential cetuximab biosimilar. Lastly, we also report herein the possible benefits of conjugating this goat derived cetuximab to an anti-cancer molecule thereby significantly increasing its potential efficacy in the current approved cancer indications.

## Material and Methods

### Cetuximab

Cetuximab was expressed in the milk of lactating goats and purified as previously described (10). Commercially sourced cetuximab was obtained from Onelink (NZ) Ltd. This material was produced by the ImClone Systems Incorporated as previously described (11–13).

### Cell culture

The melanoma cells NZM37 and NZM40 were chosen from a panel of primary melanoma cell lines that were generated from biopsies of metastatic melanoma samples from patients presenting at clinics in Auckland, New Zealand as previously described (14–17). The cells were maintained in α-modified minimal essential medium (MEM-α) supplemented with antibiotics (100 U/mL penicillin, 100 μg/mL streptomycin, and amphotericin B 0.25 μg/mL; GIBCO Life Technologies), ITS (5 μg/mL insulin, 5 μg/mL transferrin and 5 ng/mL sodium selenite; Roche Diagnostics GmbH), and 5 % fetal bovine serum (FBS, HyClone). The colorectal cancer cell line HT29 was obtained from ATCC and maintained in MEM-α supplemented with penicillin (100 U/mL), streptomycin (100 μg/mL), and 5 % fetal bovine serum (FBS).

### Western blotting

Protein concentration of total cell lysate was quantified by BCA assay. For western blotting 40 μg of protein samples were subjected to 10% in house SDS-polyacrylamide gel electrophoresis (SDS-PAGE) and transferred onto nitrocellulose membranes (Millipore). The membranes were blocked with 3% BSA in TBS containing 0.01 % Tween-20 for 1 h at room temperature and then incubated with specific primary antibodies at 4 °C overnight in the blocking buffer. After TBST washing, membranes were incubated with secondary antibody at room temperature for 1 h in the blocking buffer. Detection of specific protein expressions were performed by Clarity Western ECL blotting substrates with Bio-Rad ChemiDoc MP imaging system. Antibodies used for immunoblotting are as follows: total-EGFR (Cell Signaling Technology #2232, 1:1000), phospho-EGFR (Cell Signaling Technology #2234, 1:1000), β-Actin (Sigma #A1978, 1:2000), total-Akt (Cell Signaling #9272, 1:1000), and phospho-Akt (Cell Signaling Technology #9271, 1:1000).

### Construction of CRISPR-mediated EGFR knock out stable cell lines

A pair of guide RNA targeting human EGFR was cloned into pSpCas9(BB)-2A-GFP (px458) plasmid vector (Addgene plasmid #48137) (Addgene, Cambridge, MA) following the depositor’s protocol. The sequences are: Forward: 5’-CACCGGGAGCAGCGATGCGACCCTC-3’ and Reverse: 3’-CCCTCGTCGCTACGCTGGGAGCAAA-5’. Melanoma cell lines NZM37, NZM40 and colorectal cancer cell line HT29 were transfected with pX458-gRNA using lipofectamine 3000 (Life Technology) according to the manufacturer’s instructions. 24 h after transfection, GFP positive cells were sorted by FACSAria II SORP cell sorter and seeded into 96 well plates with single cell per well. Expression of EGFR in expanded colonies was detected by immunoblotting.

### Cell viability assay

Cells were seeded in 96-well plates (5,000 cells/well). After 24 h, cells were incubated with different drugs accordingly with a range of concentrations (from 0.1 to 100 *μ*M) for 72 h. Cell viability was determined using the sulforhodamine B (SRB) assay as previously described (18). Results were plotted as percent of vehicle control from at least two independent experiments conducted in triplicate. Growth curves were analyzed by nonlinear regression using GraphPad Prism V6.0 software (GraphPad Software, San Diego, CA).

### Synthesis and analysis of MMAE and Cy5 anti-EGFR antibody conjugates

Commercial cetuximab (ImClone, 5 mg/ml) and gCetuximab (6.32 mg/ml) were diluted to 2 mg/ml with sterile saline. Diluted cetuximab solutions (2 mg) were treated with 100 *μ*l Bicine buffer (1M, pH8.26) and 10 *μ*l diethylenetriaminepentaacetic acid (DTPA) (100 mM, pH7.0). Then antibodies were reduced by 4 equivalents of tris (carboxyethyl) phosphine (TCEP) at 37 °C for 2 h. After cooling down to room temperature, 4 equivalents of MC-VC-MMAE (MedChem Express, HY-15575) were added and incubated for 30 min. Reaction mixtures were gel-filtered through Sephadex G-25 (Sigma G2580-10G) and eluted by PBS (19). Subsequently, the conjugated cetuximab-MMAE mixtures were concentrated by centrifugal concentrator (30 kDa MWCO, Sigma). Before labelling the conjugated cetuximab-MMAE with Cy5-maleimide, 50 mM DTT was added to reduce the remaining disulphide bonds (20). After 30 min incubation, 2 equivalents of Cy5-maleimide (Abcam ab146489) was added and incubated 30 min at room temperature. Following buffer exchange through Sephadex G-25, the Cy5-cetuximab-MMAE conjugates were concentrated by centrifugal concentrators. Cy5-cetuximab-MMAE conjugates were analysed by hydrophobic interaction chromatography-HPLC using an TSKgel Ether-5PW column (Tosoh Biosciences). Antibody-drug isomers were separated from unconjugated cetuximab by the HPLC method using a linear gradient from 100 % high salt concentration buffer A (0.05 M sodium phosphate buffer, pH7.0; 2 M ammonium sulphate) to 100 % buffer B (80 % v/v 0.05 M sodium phosphate buffer, pH7.0, 20 % v/v 2-propanol) in 45 min (21). The flow rate was set at 200 *μ*l/min. The concentrations of antibodies were measured at 280 nm by Nanodrop spectrophotometer.

### Cetuximab Half-life Assessment

Female CD-1 mice, 5-6 weeks of age, were obtained and maintained in Vernon Janson Unit, University of Auckland. All studies were performed in accordance with University of Auckland animal ethics and animal welfare. Cetuximab was administrated by intraperitoneal injection (*ip*) with 12.5 mg/kg dose. Blood samples were collected at 0, 1, 3, 6, 24, 48, 72, 96, and 120 h by cardiac puncture, following a single intraperitoneal injection (*ip*) according to previous publications (22). The blood samples were centrifuged at 2000xg for 15 min at 4 °C. Plasma aliquots were stored at −80°C until analysis by an ELISA assay. The ELISA assay was performed according to the manufacturer’s instructions (ImmunoGuide IG-AB112). Briefly, the ELISA assay is based on a cetuximab-specific mouse monoclonal antibody pre-coated onto microtiter plates to capture cetuximab in mouse plasma. The captured cetuximab from plasma was then detected by a horseradish peroxidase (HRP)-conjugated anti-human IgG monoclonal antibody binding to cetuximab Fc fragment. After addition of chromogen-substrate, the colour developed is proportional to the amount of cetuximab in the sample or standard. In order to exclude the affinity differences between commercial and goat-produced cetuximab to mouse anti-cetuximab antibody coating the ELISA plate, a standard curve with goat-produced cetuximab was also included for data analysis.

### Xenograft mouse model

Colorectal cancer cells HT29 (5 × 10^6^) were injected subcutaneously into the right side of the NIH-III immunocompromised nude mice at 5-6 weeks of age. Tumour volume was calculated every 3 days with a caliper using the following formula: (π × length × width^2^) / 6, where length represents the largest tumour diameter and width represents the perpendicular tumour diameter. When the average tumour volume reached 100 mm^3^, mice were dosed every 3 days with commercial cetuximab, goat-produced cetuximab or vehicle control by intraperitoneal injection with the dosage of 10 mg/kg. The data were analysed using a two-way analysis of variance (ANOVA) model.

### Immunohistochemistry

After euthanization of the mice, the tumors were excised and preserved in freshly made 10 % neutral formalin buffer for 48 h. Then paraffin blocks were prepared and sectioned for immunohistochemical (IHC) staining. IHC procedures were perfomed as follows: paraffin sections were kept in 60 °C for 15 min and then rinsed in fresh xylene twice, 10 min for each time, following by 100 % ethanol wash (twice, 10 min for each time). Antigen retrieval was applied by incubating sections with 10 mM citric acid buffer at 95 °C for 30 min. After permeabilizing with TBST (0.1 % Triton X-100), the slides were blocked with TBST containing 2 % BSA and 5 % goat serum at room temprature for 1 h. Then slides were incubated with primary antibodies overnight at 4 °C. After washing with TBST the slides were incubated with a secondary antibody for 2 h at room temperature. Slides were sealed with mount media containing DAPI (Invitrogen, ProLong dimand). Antibodies used in immunohistochemical staining were as follows: Ki67 (Abcam, ab8191, 1:100), CD31 (Abcam, ab28364, 1:100).

## Results

### Effects of goat cetuximab on EGFR-dependent intracellular signaling

Colorectal cancer cell line HT29 as well as melanoma cancer cell lines NZM37 and NZM40 which have relatively high levels of endogenous EGFR expression were chosen for the following experiments (Figure 1A). CRISPR-medicated human EGFR knock out cell lines were generated and used as negative control. EGFR expression from expanded colonies of each cell line were detected by western blot (Fig. 1B, 1C and 1D). Two knockout clones of each cell line were chosen for the following experiments.

**Figure 1.**
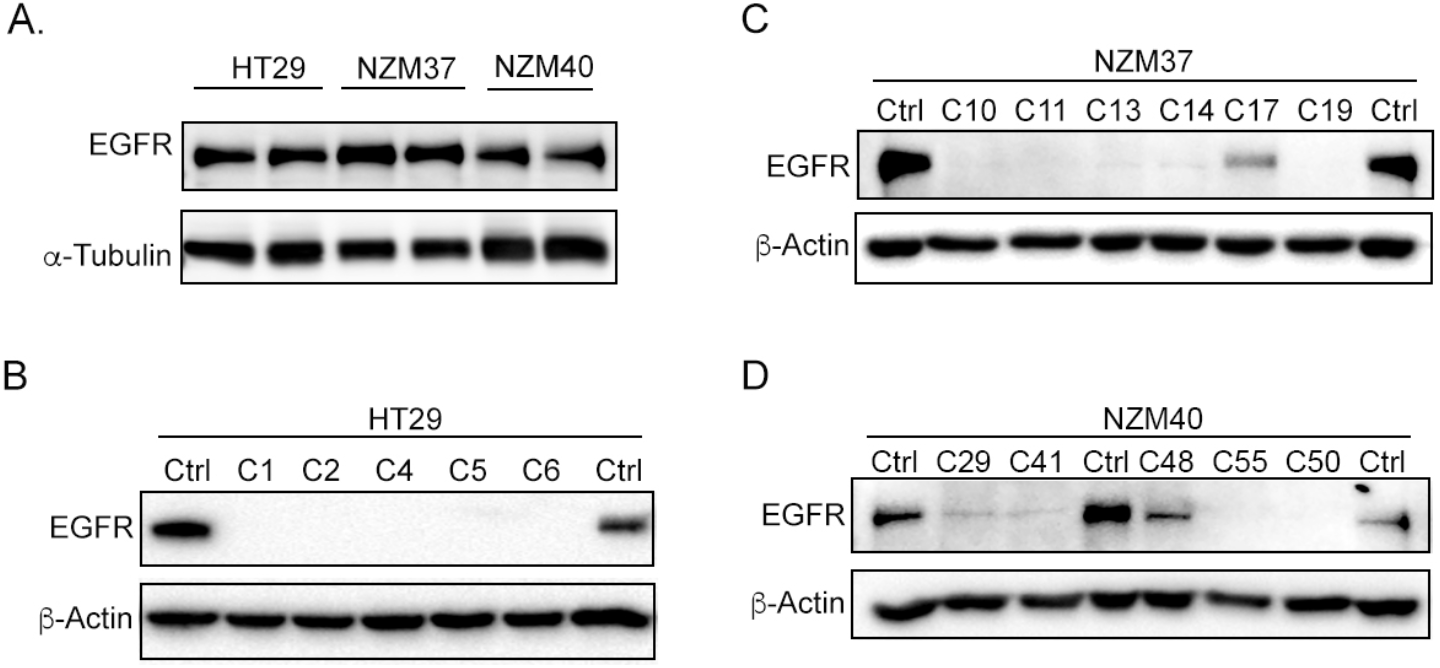
EGFR expression in colorectal and melanoma cancer cell lines. A. EGFR was endogenously expressed in both colorectal cell line HT29 and primary melanoma cancer cell lines NZM37 and NZM40. EGFR was successfully knocked out by CRISPR/Cas9 genetic editing strategy in HT29 (B), NZM37 (C), and NZM40 (D) cell lines.

In order to compare the bioactivities of gCetuximab with the commercial cetuximab, we firstly investigated the *in vitro* effects of cetuximab on the growth of EGFR expressing melanoma and colorectal cell lines using the SRB cell viability assay. After 72 h exposure, both forms of cetuximab elicited minimal growth inhibition effect on both colorectal and melanoma cells (Fig. 2A, 2B and 2C). This was consistant with previous findings that cetuximab showed limited growth inhibition on cancer cell lines growing in serum *in vitro* (23). However, It suggested gCetuximab has minimum non-specific cytotoxicity to cells and could be safely used in amimal studies. We then investigated the effects of gCetuximab on EGFR signaling by EGF ligand stimulation. Cells were serum starved for 24 h and then stimulated with EGF(100 ng/ml) for 15 min with or without cetuximab preincubation. Immunoblotting results show the complete inhibition by gCetuximab of EGFR signaling in both melanoma and colorectal cell lines (Fig. 2D, 2E and 2F), dramatically decreased phospho-EGFR and phospho-AKT levels. These results suggest the gCetuximab and commercial cetuximab have equivalent effects on human recombinant EGF stimulated EGFR signaling in melanoma and colorectal cancer cells.

**Figure 2.**
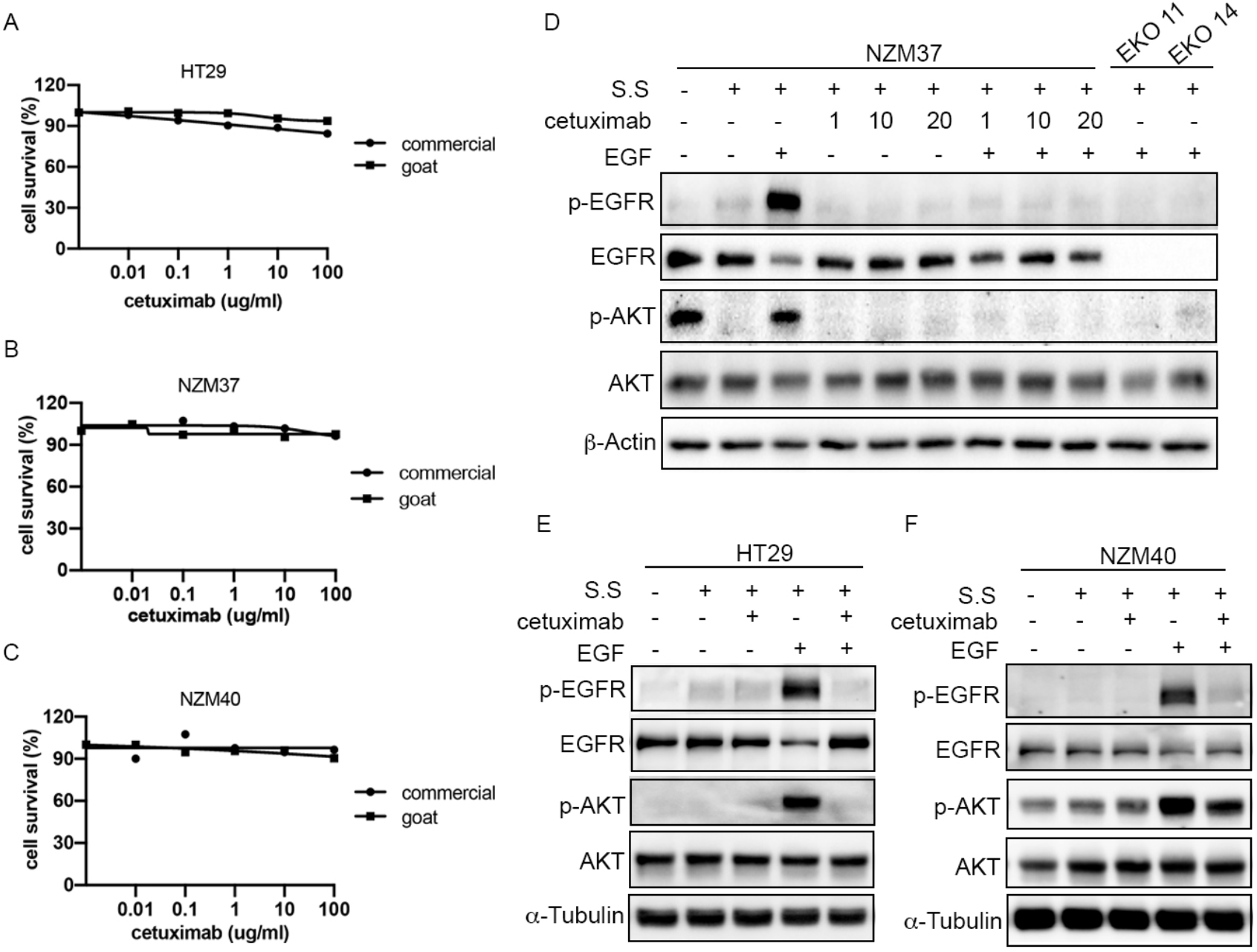
gCetuximab inhibited EGF stimulated EGFR signaling. A to C. Effects of both commercial and goat cetuximab on cell growth were tested by SRB growth assay toward HT29 (A), NZM37 (B), and NZM40 (C) cell lines; D. After overnight serum starvation, NZM37 cells were preincubated with or without gCetuximab (1, 10, 20 ug/ml), following 15min EGF (100ng/ml) stimulation. EGFR knockout clone 11 and clone 14 (EKO11, EKO14) served as negative control; E and F. HT29 and NZM40 cells were starved overnight. After preincubation with or without goat cetuximab (100ug/ml), cells were stimulated with EGF (100ng/ml) for 15min.

### Anti-tumor effects of gCetuximab on a colorectal cancer xenografted model

We next investigated the anti-tumour effect of gCetuximab using xenograft model. Both gCetuximab and commercial cetuximab significantly inhibited the growth of HT29 xenograft tumours (Figure 3A). There was no body weight loss when animals were treated with either drug indicating no obvious *in vivo* toxicity. This suggests that the similar antitumour efficacy of the gCetuximab and commercial product (Figure 3A, 3D).

**Figure 3.**
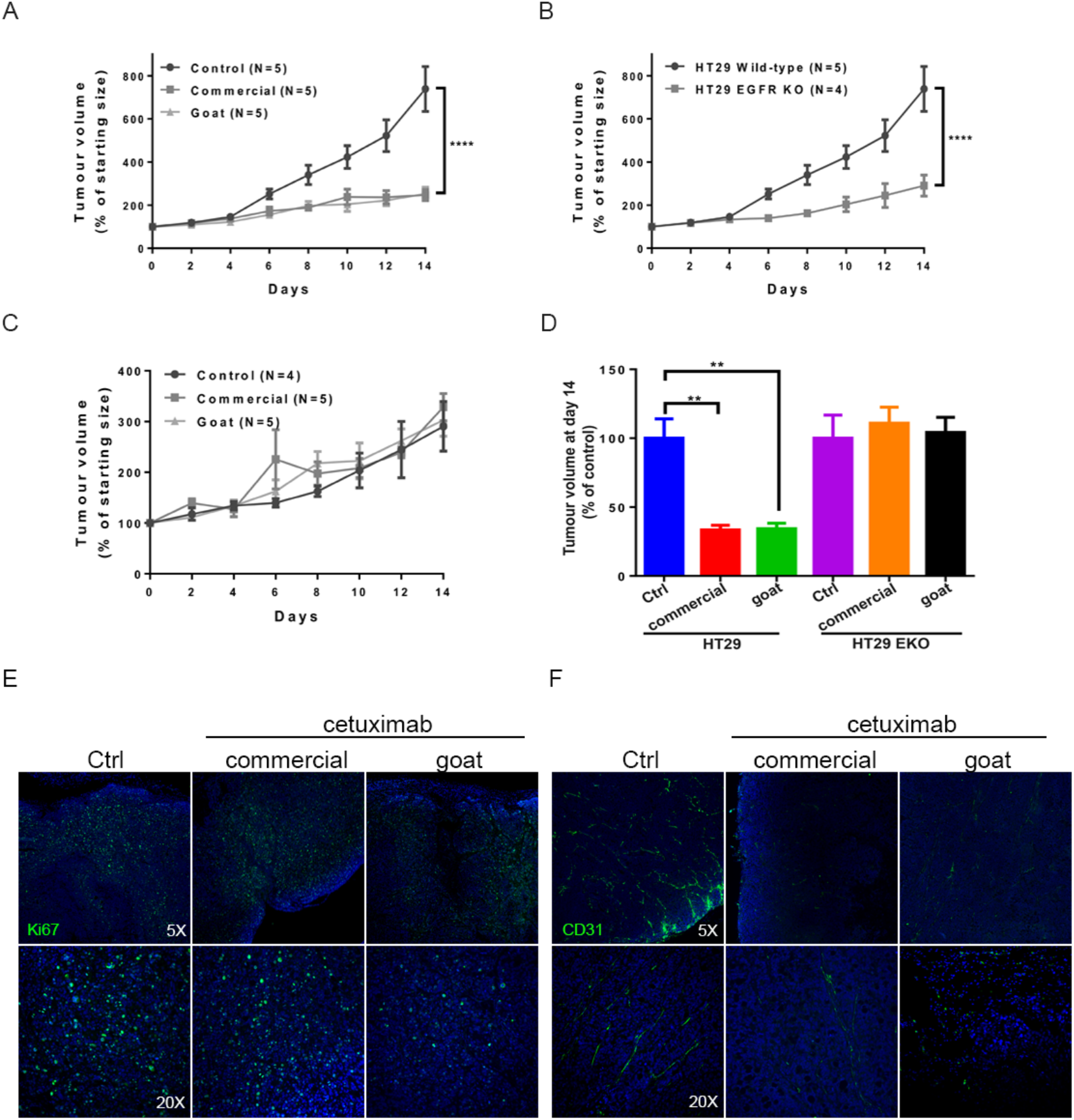
Antitumour effect of cetuximab in xenograft mice model. Xenografts were established in female NIH-III mice by subcutaneous injection of EGFR knockout or parental HT29 cells. Commercial or gCetuximab (10 mg/kg) was administered by intraperitoneal injection every 3 days. Tumour size was measured every 2 days (*n*=6). A. Effects of cetuximab on HT29 tumour growth; B. Tumour volume of EGFR knock out and parental HT29 cell lines. C. Effects of cetuximab on EGFR knock out HT29 tumour volume; D. Quantification of tumour volume at day 14. Data was analysed by t-test (*p*<0.01); E and F. Immunohistochemical (IHC) analysis of tumour proliferation maker Ki67 and microvessel marker CD31.

In order to confirm the tumor growth inhibition effect was due to the specificity of gCetuximab binding to EFGR, studies were done with the EGFR knock out cell line. When transplanted into mice, EGFR knock out HT29 tumours showed delayed tumour growth compared to HT29 parental tumours with its level being similar to that of the wild type HT29 xenografts that were treated with cetuximab (Figure 3B). When these tumors were treated with either of the antibodies there was no effect on the growth of EGFR knock out HT29 xenograft tumours (Figure 3C). Together this shows that the drugs were specifically targeting EGFR on HT29 cancer cells in these xenograft models.

Immunohistochemical (IHC) analysis was performed to investigate the mitotic index of cetuximab treated tumours, based on the proliferation marker Ki67. In addition, microvessel density of cetuximab-treated xenograft tumours was also assessed using CD31 as a marker. The results showed a reduction of Ki67-positive staining in tumours following cetuximab treatment relative to the untreated control (Figure 3E). Similarly, the microvascular density (CD31-positive cells) was also significantly decreased after cetuximab treatment compared to untreated controls (Figure 3F). For both Ki67 and CD31 there was no significant difference between commercial and gCetuximab treatments. Taken together, these data again suggested that gCetuximab could effectively inhibit tumor growth via EGFR signaling *in vivo*.

### Effects of toxin conjugated goat and cell culture produced cetuximab on colorectal cancer cells

Antibody-drug conjugates (ADCs) for cancer therapy has been attracting significant attention over the past few years due to its precision in cancer target therapy. Monomethyl auristatin E (MMAE), a potent antimitotic drug, cannot be used as a drug itself due to its toxicity. Instead, it can be linked to monoclonal antibodies to form ADCs for targeted cancer therapy. In this study, we tested the ability of gCetuximab to be linked with MMAE and investigated the effectiveness of gCetuximab-MMAE conjugates at targeting toxins to tumors by comparing to commercial cetuximab product.

By using a reducing reagent, MMAE was successfully linked to both the gCetuximab and commercial cell culture produced cetuximab. To test if these two ADCs can deliver MMAE specifically to tumour cells, the ADCs were labelled with Cy5 for tracking. EGFR expressing colorectal cancer cell line HT29 was treated with gCetuximab ADC or control cetuximab ADC and analyzed for Cy5 fluorescence. After 30 min, both ADCs were specifically localized at the cell surface and were gradually degraded within 24 h concurrent with EGF receptor internalization (Figure 4A). We then investigated the cell killing effect of both ADC conjugates by *in vitro* SRB cell growth assay. Both ADCs were more effective than cetuximab alone at killing HT29 colorectal cancer cells (Figure 4B, 4C). In contrast, both ADCs were unable to kill HT29 EGFR knockout cells (Figure 4D, 4E).

**Figure 4.**
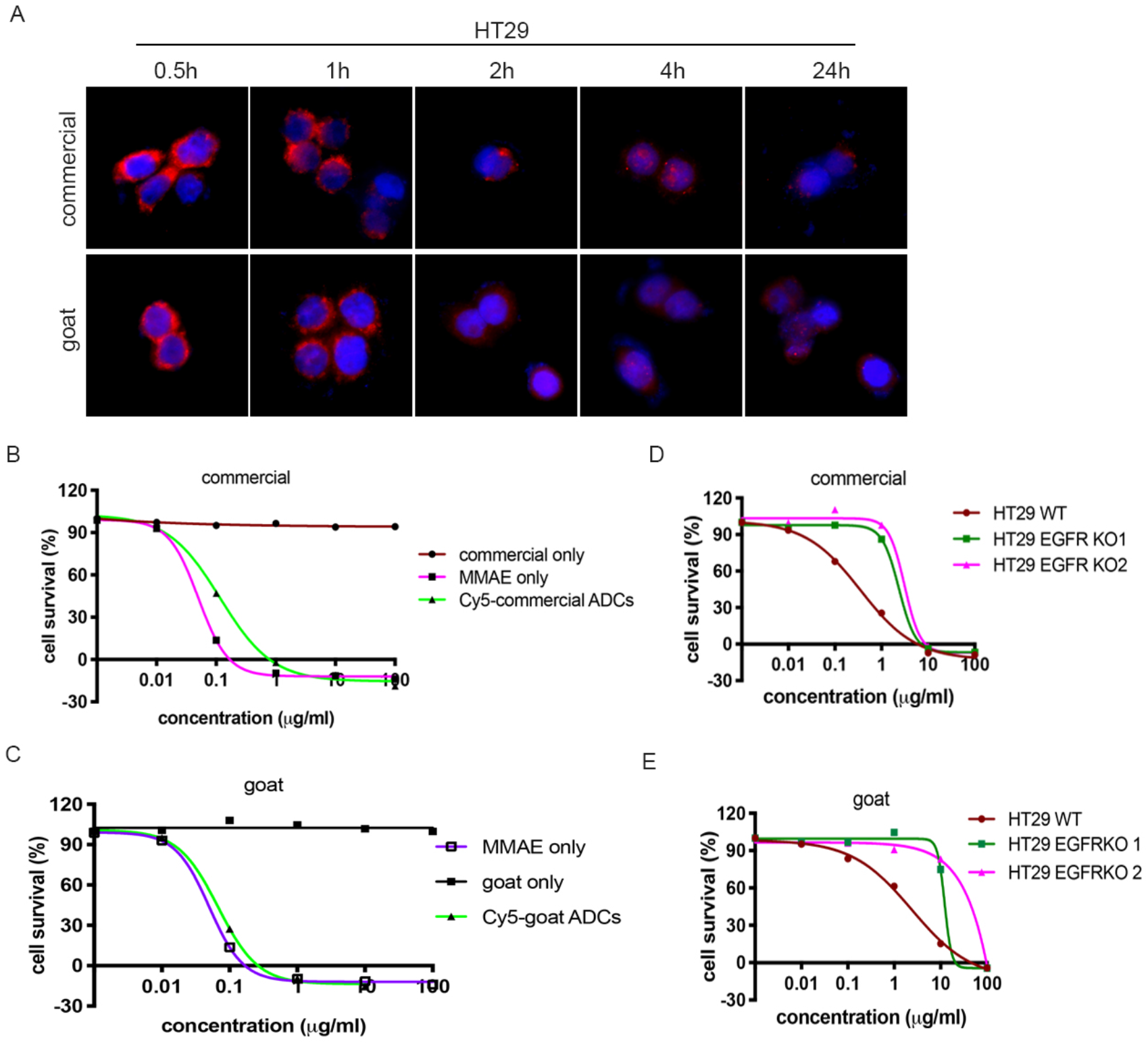
Cetuximab conjugated to toxin MMAE. A. Colorectal cancer cells HT29 were incubated with Cy5-ADCs for 0.5 h and then replaced with fresh culture media for another 1, 2, 4, and 24 h. Cy5-ADCs could specifically bind to the cell surface and gradually degraded along with EGF receptor internalization and degradation process. B and C. HT29 cells were incubated with cetuximab only, MMAE only or cy5-ADCs for 72 h. SRB cell growth assay was performed to determine cell growth inhibition by different treatments. (B. commercial; C. goat). To test the specificity of ADCs toxicity, EGFR knockout HT29 cells were also tested in SRB cell growth assay. (D. commercial; E. goat).

### In vivo half-life of gCetuximab compared to commercial cetuximab

To investigate cetuximab half-life, blood samples were collected by cardiac puncture from CD1 mice following a single dose 12.5 mg/kg of either form of cetuximab. Plasma concentration of cetuximab was quantified by ELISA assay. Cetuximab exhibited a one phase exponential decay. In order to exclude the difference of binding affinity between commercial and gCetuximab to precoated ELISA plates, we performed two sets of standard curves: gCetuximab and commercial cetuximab standard curves. There was no significant difference between these two sets of standard curves (Figure 5A, 5B). gCetuximab and commercial cetuximab plasma concentrations were calculated according to the standard curve, respectively. The maximum plasma concentration of gCetuximab was 107 μg/ml which was peaked at 6 h post-administration; while for commercial cetuximab a similar maximum plasma concentration, 108 μg/ml, was peaked at 3 h after administration (Figure 5C, 5D). The half-life (t_1/2_) of gCetuximab was calculated as being 260 h comparing to that of commercial cetuximab at 206 h (Figure 5E). There were no significant statistical differences in maximum plasma concentration and t_1/2_ between commercial and gCetuximab. It suggested similar pharmacokinetics of gCetuximab and commercial cetuximab in mice.

**Figure 5.**
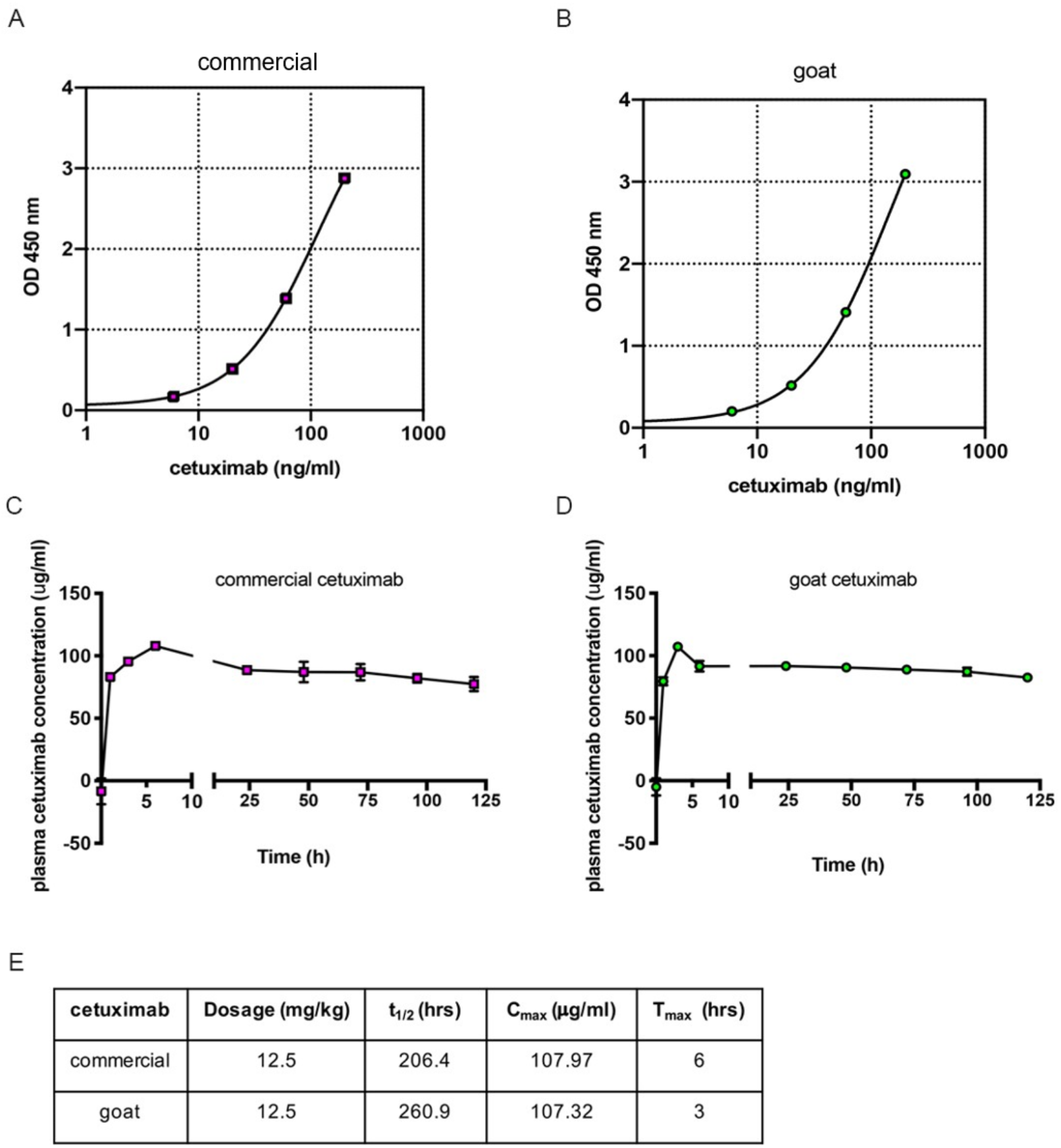
Cetuximab half-life in CD1 mice. Standard curves of ELISA assay for commercial (A) and goat (B) cetuximab. A. CD1 mice were administered with a single dose of either commercial (C) or gCetuximab (D) (12.5 mg/kg). Plasma were sampled at 0, 1, 3, 6, 24, 48, 72, 96 and 120 h (*n*=3). Cetuximab concentration in plasma samples were determined by ELISA assay. Pharmacokinetic parameters are listed in (E).

## Discussion and Conclusion

High costs of monoclonal anti-cancer antibody therapies restricts their overall routine use in the clinic. We have previously described a transgenic animal production system for the monoclonal antibody cetuximab in which high levels of the therapeutic proteins are expressed in the mammary gland of lactating goats (10). In this study, we investigated antitumour effects of goat-produced cetuximab by a range of biological assays both *in vitro* and *in vivo*. Our results provide evidence that the cetuximab produced in and purified from goat milk is as equally effective in targeting and inhibiting the EGF receptor as is the cetuximab produced by traditional cell culture based methods. Despite the differences in the production systems, the goat-produced cetuximab showed no obvious difference in its toxicity profile in mice as assessed by weight loss. We have previously reported on the transgenic goat derived cetuximab relative to an expected increased safety profile due to a lack of potential immunogenicity (i.e. no α-gal linkages) as is seen with the innovator and mammalian cell culture derived cetuximab product. Additionally, we have previously reported on the potential for increased efficacy from the transgenic goat milk derived cetuximab based on its increased antibody dependant cellular toxicity (ADCC) profile (10). We now report that, compared to the commercially available cetuximab, the goat derived cetuximab is equally suited as a antibody for attaching a toxin to create an ADC form of cetuximab. We show that it can target the toxin MMAE to HT29 cells and that this resulted in killing of the cells. This shows the gCetuximab is a good vehicle for treatment regimes based on antibody drug conjugates.

Taken together, this work confirms that the goat-produced cetuximab is as effective as the current commercial product and therefore suitable as a candidate for clinical biosimilar development. Further, this provides continuing support for the goat milk production system as a commercial and proven (mulitple agency approvals worldwide) platform for cost effective human recombinant protein therpatuic production and now to be applied to potetnial biosimilars going forward.

## Acknowledgements

Funding was provided by the Ministry of Business Innovation and Employment in New Zealand (contract ID is C10X1504), AgResearch and from the Maurice Wilkins Centre for Molecular Biodiscovery. The authors wish to thank Lynn Peters for the purification work and Dan Pollock, Harry Meade and Lihow Chen for reviewing the draft and also Christina Buchanan for assistance with HPLC analysis. We thank Dr Moana Tercel for advice on developing ADCs.

